# StrPhaser constructs tandem repeat alleles from VCF data

**DOI:** 10.1101/2025.01.22.634325

**Authors:** Xuewen Wang, Jonathan King, Huang Meng, Michael D. Coble, August E. Woerner

## Abstract

Variant calling is a ubiquitous genomic technique that underpins many scientific disciplines. From a computational perspective, variant calling is a form of logical compression; neglecting large variation, a person’s genome can be losslessly described as a set of differences (SNP and small InDel alleles) relative to the reference sequence. Another common genomic technique is haplotype phasing, wherein alleles are partitioned into their paternal and maternal components (as haplotypes). Some classes of alleles are more difficult to describe than others, e.g., short tandem repeats (STRs). STRs serve as a critical marker for many genetic assays. However, STRs tend not to be explicitly reported in most genomic workflows. Here, we present StrPhaser, a novel algorithm that leverages phased variant calling datasets in the VCF file format to construct STR alleles. We evaluated StrPhaser on ∼10,000 STR alleles from 284 human genomes, achieving an average allele accuracy of 91%. In addition, StrPhaser better recovers longer STR alleles than competing approaches; in principle, STR alleles that are longer than the maximum read length can be characterized. This capability, combined with its user-friendly interface, speed, and generation of both STR genotypes and visualizations, makes StrPhaser a valuable tool for a wide range of genomic studies.

**Availability:** The StrPhaser is publicly available at https://github.com/XuewenWangUGA/StrPhaser.

## Introduction

DNA variants, both within and between individuals of a species, are foundational elements of genetic analyses. By far, the most common file format for genetic variants is the variant call format (VCF). The VCF file format encapsulates information about single nucleotide polymorphisms (SNPs), insertions, and deletions (InDels), as well as other more complex sequence variations [1].

Tandem repeats (TRs) constitute a category of complex DNA sequence variants. TRs encompass short tandem repeats (STRs), also referred to as microsatellites, as well as other satellites (e.g., minisatellites). STRs are characterized by some (often variable) number of tandem repeats [2, 3]. STRs, with a repeat unit of 1<n≤6 base pairs, are ubiquitously dispersed (∼1%–3%) across most genomes [4-7]. TRs tend to be highly polymorphic, predominantly due to unit expansion and contraction. While not typically characterized as such, TRs can be described as a sequence of (predominantly InDel) variants. In a phased VCF, variants inherited from either parent are presented separately; thus, TR information can be gleaned from a VCF file. However, TR alleles are not (directly) reported in VCFs by widely used variant call tools such as GATK [8, 9] and DeepVariant (Poplin et al., 2018), despite the development of several TR specialized *de novo* calling tools, specifically for STRs [2, 4, 10].

Calling STRs from whole genome sequencing is a challenging computational problem. Most approaches would only consider reads that span the entirety of an STR region [2, 4, 10-13]. As the read length approaches the length of the STR, STR characterization becomes technically challenging. As an example, if a STR allele is half of the read length, only ∼1/2 of the reads from a shotgun sequencing assay will fully span the repeat. Variant calling, however, serves to decompose a STR into heterogeneous parts; thus, it is still possible to characterize (compound and complex) STRs even if the reads only partially overlap the region of interest.

A modern genomic tool for mining sequence-based STR alleles from the genome-wide variant data is vital to STR applications, like forensic genetics. Here, we have introduced a novel algorithm to construct STR alleles from phased VCF data and implemented it in a new bioinformatics tool, StrPhaser. The method can work for any kind of TRs, including STRs. StrPhaser was assessed on a unique dataset of 289 individuals sequenced on two competing sequencing technologies.

## Results

### The outline of the StrPhaser algorithms

StrPhaser requires the genomic coordinates of the targeted STR or TR regions, the reference sequence in Fasta format, and a VCF file that has been genotyped, phased, and is devoid of missing data. The VCF data are evaluated for each TR region. Each phased genotype is categorized by its type—SNP, insertion, or deletion. Note that multibase SNPs (MNPs) are treated as multiple SNPs, and only alternative genotypes are considered. StrPhaser reconstructs each TR allele based on the phased variant annotations (**Figure 1**).

**Figure 1.**
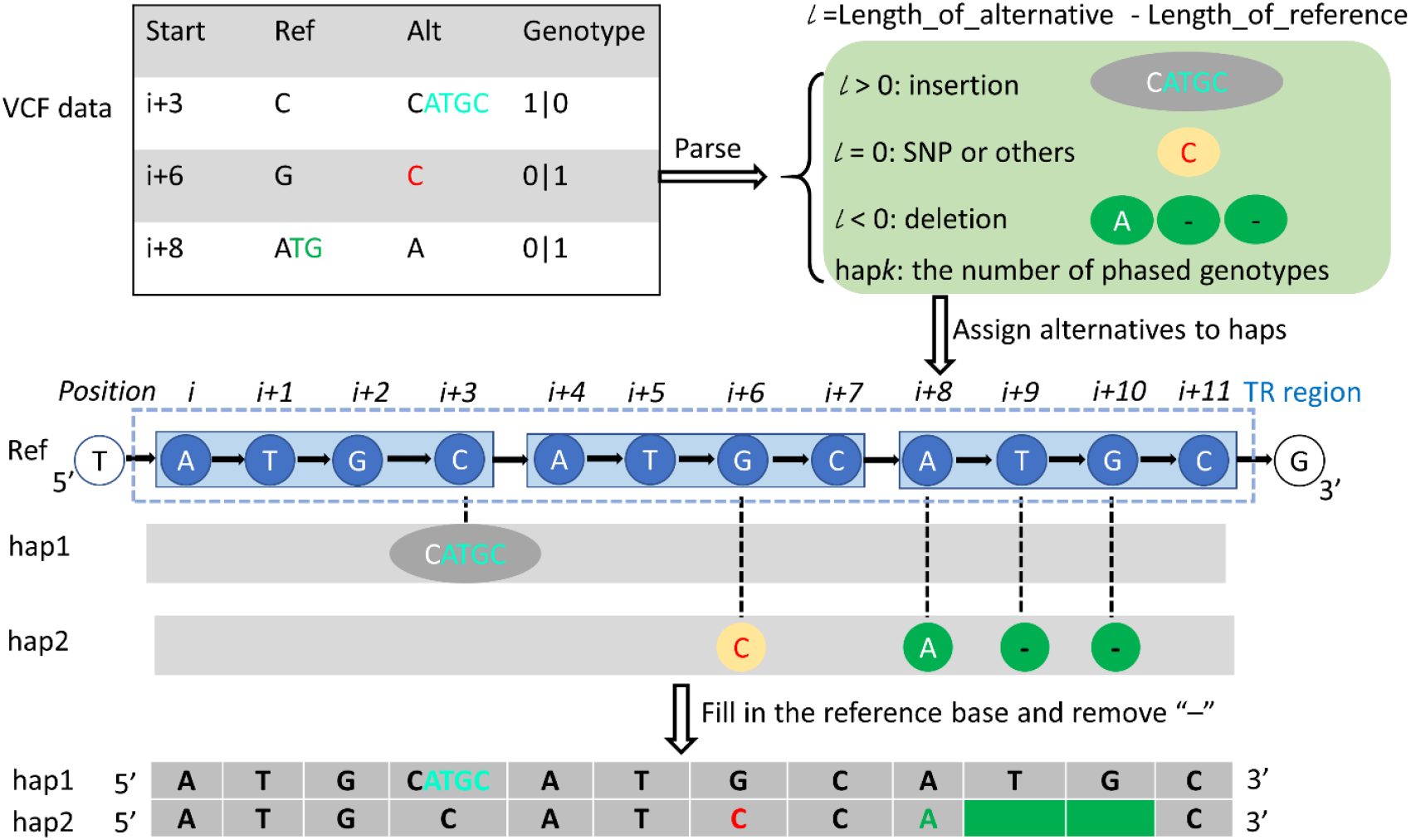
An illustration of the algorithms in StrPhaser. Tandem repeat (TR) alleles span some number of single nucleotide polymorphisms (SNPs), insertions, and deletions of variant annotations (VCF data). SNPs, insertions, and deletions correspond to substitution, insertion, and deletion operations, respectively, applied to the reference sequence. Each operation has a corresponding position (e.g., i + 3), as well as the allele(s) which are derived from the Ref, and Alt categories and the sample genotype.

Initially during the parsing step, VCF data are partitioned to the vicinity of each targeted TR region. Each phased genotype in the VCF is parsed; as variant calls may occasionally overlap, only alternative alleles are considered. This process involves defining the length (*l*) as the difference in nucleotide count between an alternative genotype and the reference genotype at a VCF site (**Figure 1**). If *l* > 0, the allele is classified as an insertion and parsed accordingly (e.g., **Figure 1**, CATGC). If *l* = 0, the alternative is identified as an SNP or equivalent and is parsed as one exact alternative base as in the VCF (e.g., C). If *l* < 0, the alternative is a deletion, where the alternative base(s) will be parsed as the first base of the genotype in the reference and “-” for all deleted base(s) (e.g., A--). The number of repeated “-” symbols is calculated as the length of the reference bases minus one.

In the step of assigning variants to haplotype, the reference coordinates at a given TR region and their corresponding bases will be stored in two rows of a table in the order of 5′ to 3′ of the reference DNA sequence, where each positional column holds one nucleotide. This is a simplified data structure, where each node is a base and each edge is from the previous base to the next base. These two rows hold the reference allele information. Then, additional rows/paths will be added to the table/graph to represent the haplotypes, e.g., hap1, and hap2 for two haplotypes (**Figure 1**). The mutated node at the given coordinate will be assigned with the parsed bases from the previous step. For insertion and SNP alleles, alternative bases will be directly assigned to the corresponding node, e.g., CATGC at position *i+3* of hap1 (**Figure 1**). For deletions, each character of the parsed bases will be assigned to a node, respectively, starting from the deletion starting position, e.g., positions *i+8, i+9*, and *i+10*, respectively (**Figure 1**). After parsing variants, all intervening bases are treated as the reference sequence.

Given the complexity of TRs and the ambiguity of where a TR begins and ends, StrPhaser provides an option to characterize (and include) flanking variation. With this option, bases adjacent to the targeted region are automatically included. Following the initial construction of the allele, extended bases at the end of the allele are trimmed off to restore to the user-defined region prior to reporting the final allele. Manual inspection often reveals that more accurate alleles can be identified with this option.

### StrPhaser input and output

StrPhaser requires a VCF file and a configuration file that describes the TRs. The VCF file supplies variant information, which includes InDels and SNPs of TR regions. The TR configuration file, formatted as tab-separated plain text in BED file format, provides the targeted TR coordinates and the corresponding TR site names (Table 1). An exemplary TR configuration file for three forensic Combined DNA Index System (CODIS) STRs is presented in **Table 1**. Additional arguments, e.g., computing threads, are also available but optional.

**Table 1.** A configuration file of targeted STR sites in the human genome.

| Chrom | Start | End | Name |
| --- | --- | --- | --- |
| chr1 | 230769616 | 230769683 | D1S1656 |
| chr2 | 1489653 | 1489684 | TPOX |
| chr8 | 124894865 | 124894916 | D8S1179 |

*Chrom, Start, and End represent chromosome name, the 0-based start coordinate, and the 1-based end coordinate in the human genome reference hg38, respectively, while the Name is the description for the locus.*

StrPhaser is written in the Java computing language with multiprocessing support. It offers a user-friendly, command-line interface and is compatible with various operating systems including Windows 10, Linux Ubuntu 20.04, Centos 8, and Mac OS 10 or later. StrPhaser is fast. During internal testing, the software was able to construct alleles for all 20 CODIS core STR sites from the 1000 Genomes Project (1KG) phased VCFs in approximately one second. This was achieved using two parallel computing threads on a Linux machine equipped with an AMD EPYC 7713 Processor.

### STR allele construction and comparison for core CODIS forensic sites in 289 humans

StrPhaser was used to construct the alleles for the 20 core CODIS STR loci. Phased VCF data [3] from the 1KG project (2×150bp, ∼30×) were compared to 289 overlapping individuals sequenced on a MiSeq FGx (351bp for the long fragment and 31 bp for the short fragment, targeted sequencing) and characterized by the tool UAS [14]. To ensure that STRs were genotyped consistently, the dataset of [14] was recalled with TRcaller using locus definitions equivalent to StrPhaser (**Figure 3**). Herein, consistent data between both call sets are treated as ground truth. In total, 10,343 consistent STR alleles were obtained. StrPhaser reported 10,076 STR alleles from the 1KG dataset, of which 83.4% were correctly called (**Figure 3** A). Further analysis revealed that constructed alleles from StrPhaser had 557 (5.4%) drop-in and 479 (4.6%) drop-out alleles (See Methods; Supplementary Data S1). Upon manual inspection, large inconsistencies were found at four STRs: FGA, D1S1656, D3S1358, and D12S391, though the underlying etiology remains elusive (**Figure 3**B). Considering the 16 remaining loci, the average allele accuracy was 91% (**Figure 3**B).

**Figure 2.**
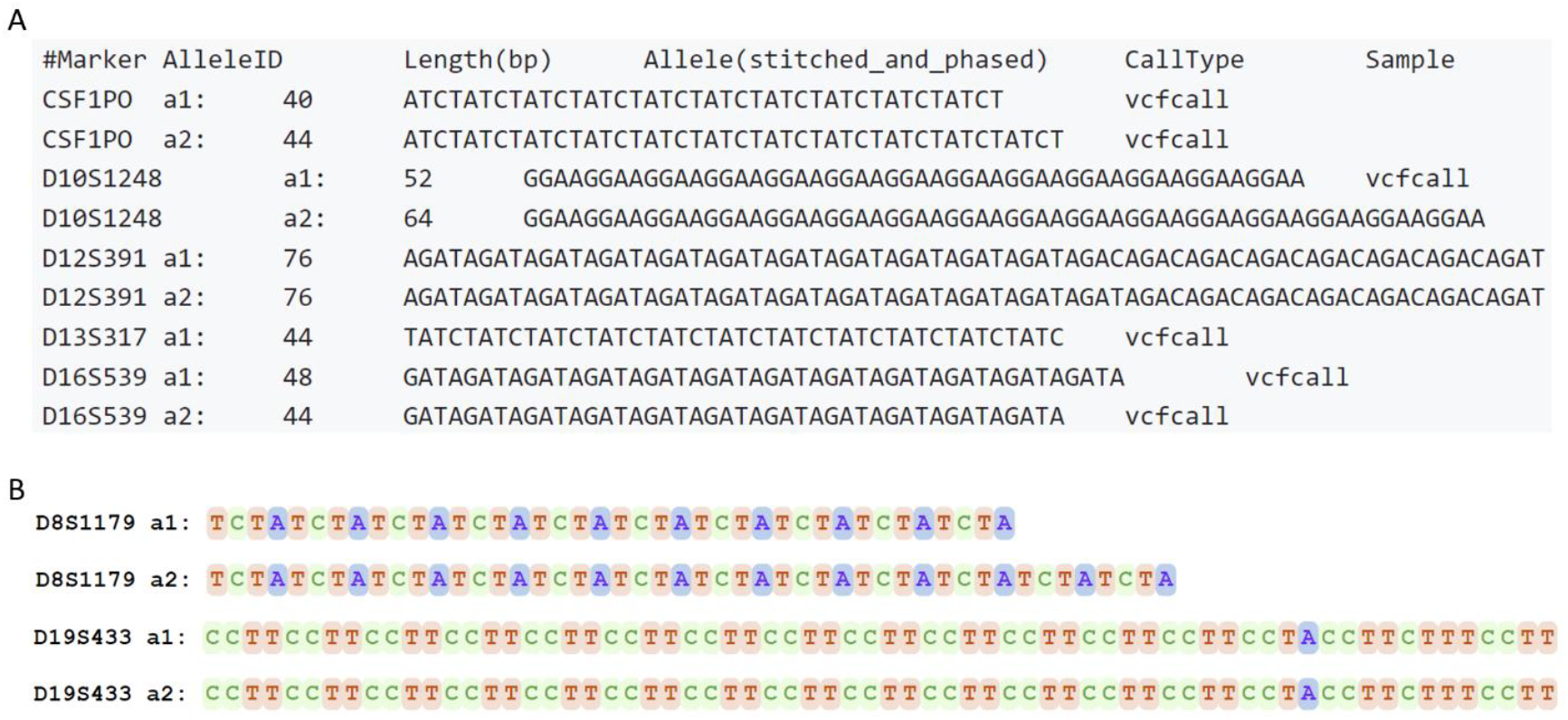
Output of StrPhaser. A. A depiction of the Tandem Repeat (TR) alleles derived from StrPhaser for five specific short TR sites. B. A color-coded representation of the allelic sequences for two selected sites shows the diversity and complexity of these sequences.

**Figure 3.**
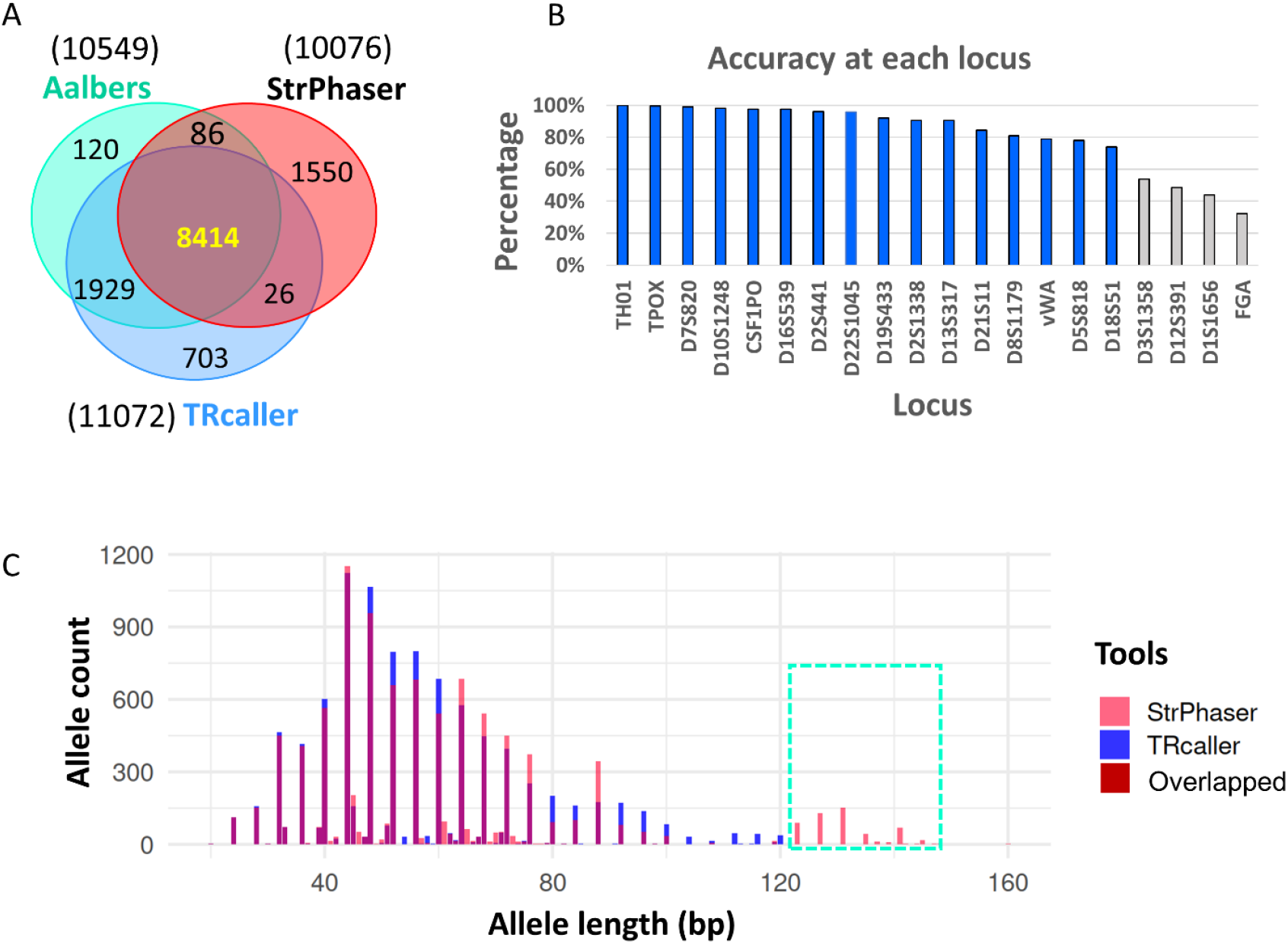
Validation and accuracy of StrPhaser constructed STR alleles. **A.** Venn diagram illustrating the overlap and uniqueness of Short Tandem Repeat (STR) alleles identified by each of the three datasets under consideration. **B**. A representation of the accuracy of constructed STR alleles for each forensic STR site. Here, the accuracy is defined as the percentage of consistent STR alleles relative to the ground truth, determined by the overlap between the Aalbers’ dataset and the results from TRcaller. The labels at the horizontal axis show the names of STR sites ordered by percentage from high to low. **C**. Comparison of allele counts versus length from TRcaller and StrPhaser, showing the long STR alleles can be called by StrPhaser only in the dashed box.

To provide a baseline, TRcaller was also evaluated on the 289 samples from 1KG. StrPhaser achieved an allele accuracy of 90.2% and a genotype accuracy of 71.0%. The lower genotype accuracy likely stems from inaccurate variant calls within the VCF dataset. To assess the ability of both tools to construct STR alleles from partially spanned reads, we compared their allele call rates across varying allele lengths. While both tools performed similarly for alleles shorter than 120 bp, StrPhaser correctly called more alleles than TRcaller, 535 versus 25, if alleles are longer than 120 bp (**Figure 3**C). Notably, these longer alleles, though less frequently covered, were recovered with 83.4% allele accuracy. This finding underscores StrPhaser’s capability to call STR alleles from limited read coverage.

### Future directions

Sample genotyping and phasing alike are probabilistic, incorporating these uncertainties will be incorporated into future versions of StrPhaser. In addition, the algorithms employed by StrPhaser assume that the accounting of variants in the VCF file is complete. i.e., all true variable sites are described in the VCF file. Further consideration of genotype refinement, imputation, and phasing may help improve STR genotyping.

## Conclusion

Variant calling and haplotype phasing are genomic standards, while the extraction of complex alleles like those found in STRs is thought to involve separate (and highly specialized) informatic strategies. StrPhaser demonstrates that STRs can be recovered from datasets encoded in the VCF file format, suggesting that variant calling may provide an additional route to characterize STRs. In addition, treating an STR as a series of variant calls permits flexibility as it removes the requirement for a read to span the entire locus, and it permits a more complete description of linkage disequilibrium between STRs and adjacent SNPs.

## Methods and materials

### Data resource

STR genotypes for the 20 core CODIS autosomal markers were drawn from the previous publication [14] STR alleles were converted to the positive strand of the human genome reference GRCh38 as necessary. Extraneous leading/trailing sequences were manually removed. In addition, raw reads (Illumina MiSeq) from the same study [14] were downloaded (NCBI Sequence Read Archive accession number PRJNA795193) and mapped to the GRCh38 reference genome (BWA, version 0.7.17-r1188) using the default settings [15]. STR alleles were called with TRcaller (version 2.0, https://github.com/XuewenWangUGA/TRcaller, parameters -c 2 -t 20 -r 0.02 -s 1) [4]. The consistent STR alleles with identical sequences for the same individual to both TRcaller and alleles from Aalbers dataset [14] were considered as the ground truth.

289 sample IDs in the Aalbers dataset [14] were also characterized as part of the 1000 Genome Project. Phased sample genotypes were retrieved (https://www.internationalgenome.org/data-portal/data-collection/30x-grch38) [3] and used as the input for StrPhaser. For these VCF datasets from the 1KG project, the genotyping was performed using GATK (version 3.5.0, McKenna, 2010); phasing was performed using Shapeit2 [16, 17]. A drop-in allele is an allele in an individual called by StrPhaser but not present in the ground truth. A drop-out allele in an individual is the one missing in the StrPhaser call but is present in the ground truth. Allele accuracy is the percentage of correct alleles identified by StrPhaser relative to the total number of ground truth alleles, as formulated below.

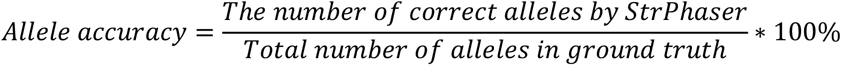

Genotype accuracy is the percentage of correct genotypes called by StrPhaser relative to the total number of ground truth genotypes.

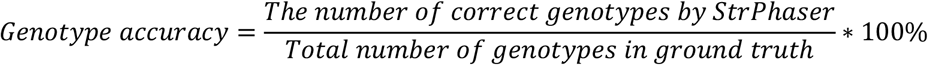

The scripts and supplementary data generated from this study are available at https://github.com/XuewenWangUGA/StrPhaser. The version 1.1 of StrPhaser is used in this study.

## Funding

This work was supported by award 15PNIJ-21-GG-04159-RESS from the National Institute of Justice and internal funds from the Center for Human Identification. The opinions, findings, and conclusions or recommendations expressed are those of the authors and do not necessarily reflect those of the U.S. National Institute of Justice.

## Contributions

A.W. designed the study. A.W. and M.C. supervised the project. X.W. and A.W. developed the algorithms and software. X.W. validated the software and analyzed the data. X.W. and H.M. tested the software. X.W. drafted the initial manuscript. J.K., A.W., and M.C. revised the manuscript. All authors approved the final version.

## Confliction of interest statement

The authors declare no conflicts of interest.

## Notes

https://github.com/XuewenWangUGA/StrPhaser

### Competing Interest Statement

The authors have declared no competing interest.

